# The neuroinflammatory interleukin-12 signaling pathway drives Alzheimer’s disease-like pathology by perturbing oligodendrocyte survival and neuronal homeostasis

**DOI:** 10.1101/2021.04.25.441313

**Authors:** Shirin Schneeberger, Seung Joon Kim, Pascale Eede, Anastasiya Boltengagen, Caroline Braeuning, Myrto Andreadou, Burkhard Becher, Nikos Karaiskos, Christine Kocks, Nikolaus Rajewsky, Frank L. Heppner

## Abstract

Alzheimer’s disease (AD) is characterized by deposition of pathological amyloid-β (Aβ) and tau protein aggregates and involves chronic neuroinflammation, ultimately leading to neurodegeneration and cognitive decline. Central in AD-related neuroinflammation is the proinflammatory interleukin-12 (IL-12)/IL-23 signaling pathway whose inhibition has been shown to attenuate pathology and cognitive defects in AD-like mice. In order to explore which cell types are involved in this neuroinflammatory cascade, we used single-nuclei RNA sequencing in AD-like APPPS1 mice lacking or harboring IL-12/IL-23 signaling. We found *Il12b* transcripts encoding the common p40 subunit of IL-12/IL-23 signaling to be expressed preferentially, but not exclusively, in microglia in an AD-specific manner. In contrast, transcripts for the other subunits of the IL-12 signaling pathway were expressed constitutively in neurons and oligodendrocytes irrespective of AD pathology, while transcripts for IL-23 were almost undetectable. Notably, genetic ablation of IL-12/IL-23 signaling did not affect the inflammatory gene expression profile of the AD-specific disease associated microglia (DAM), but reversed the loss of mature myelin-producing oligodendrocytes and alterations in neuronal homeostasis in APPPS1 mice. Taken together, our results reveal that IL-12, but not IL-23 is the main driver of AD-specific IL-12/IL-23 neuroinflammation, which alters neuronal and oligodendrocyte functions. Given that drugs targeting IL-12 already exist, our data may foster first clinical trials in AD subjects using this novel neuroimmune target.

## Introduction

An estimated 45 million people worldwide are suffering from Alzheimer’s disease (AD)^1^, the most prevalent cause of dementia. This detrimental neurodegenerative disorder is characterized by progressive cognitive and functional impairment ultimately leading to memory loss. Pathological hallmarks of AD are the faulty aggregation and deposition of amyloid-β (Aβ) and tau proteins as well as pronounced neuroinflammation, which escalate with disease development. This process is primarily driven by the brain’s intrinsic myeloid cells - the microglia^2^. In AD, microglia acquire an altered phenotype, aggregate next to Aβ plaques^3^ and secrete soluble inflammatory factors^4^. Neuroinflammation is increasingly recognized as a driver of disease progression and thus an opportunity for novel therapeutical interventions^5,6^. Yet other glial cells, such as oligodendrocytes, are also emerging as active contributors to neuroinflammation and have been shown to be perturbed in the course of AD^7,8,9^.

A key inflammatory pathway in AD pathology is interleukin (IL)-12 and IL-23 signaling. IL-12 levels are increased in brain tissue and the cerebral spinal fluid (CSF) of AD and mild cognitive impairment (MCI) patients^10^. IL-12 and IL-23 are heterodimers comprising the subunits p35 and p19, while sharing the common subunit p40 (*Il12b*). p40 can also form homodimers, which binds either as homo- or heterodimer to the IL-12 receptor subunit β1 (*Il12rb1*) (Fig. 1a). IL-12-specific signaling is executed when – in combination with p40 actions - the IL-12 subunit p35 (*Il12a*) binds to IL-12 receptor subunit β2 (*Il12rb2*), while IL-23-specific signaling is induced in concert with p40 by the IL-23 subunit p19 (*Il23a*) binding to the IL-23 receptor (*Il23r*)^11,12^. Simultaneous inhibition of both IL-12 and IL-23 signaling by genetic removal of the shared p40 subunit, or by pharmacologically blocking p40 using specific antibodies resulted in a substantial reduction in AD-related pathology in transgenic mouse models of amyloidosis^13,14^.

**Figure 1:**
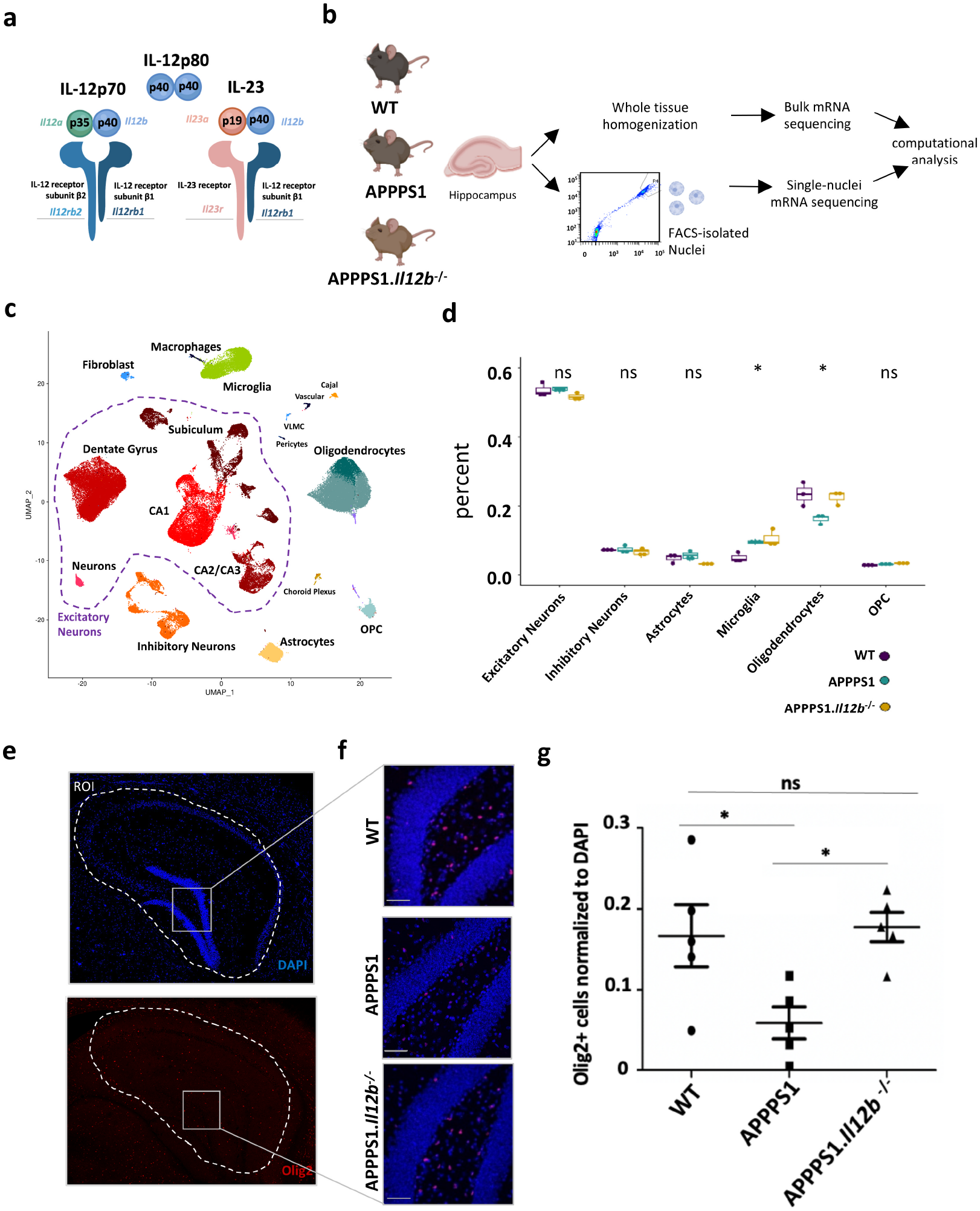
IL-12/Il-23 signaling causes loss of hippocampal oligodendrocytes in a mouse model of Alzheimer’s disease. **a)** p40 can form monodimer as IL12-p80 or heterodimer as IL-12p70 consisting of p35 and p40. IL12p70 binds to the dimerized IL-12Rβ2 and IL-12Rβ1. IL-23, consisting of p19 and p40, binds to the receptor subunits IL-23R and IL-12Rβ1. The genes that encode the respective protein subunits are shown in matched color. **b)** Experimental outline. Nuclei were isolated from individual hippocampi (n=3 per genotype) from 250 day old animals, purified by FACS and used for single-nucleus RNAseq. Bulk RNAseq libraries were prepared from RNA isolated from intact hippocampi (n=3 per genotype). Bulk RNAseq data has been generated from n=3 per genotype. If not state otherwise, all following figures reflect data from snRNA-seq. **c)** UMAP plot showing 37 hippocampal cell clusters representing combined snRNAseq data from three biological replicates per genotype. **d)** Distribution of cellular proportions across all three genotypes analyzed by one-way ANOVA. Each dot represents one biological replicate. * p ≤ 0,05, ns = not significant **e)** Using DAPI, the hippocampal outline was set as ROI for the Olig2 stained brain sections. **f)** Zoomed-in representative images of WT, APPPS1 and APPPS1.*Il12b* ^-/-^ **g)** Quantification of Olig2 positive cells normalized to DAPI positive cells in hippocampal regions. n= 5 per genotype with 3-6 sections per animal. * p ≤ 0.05, ns = not significant. Analyzed by oneway ANOVA with Tukey’s multiple-comparisons test. Each symbol represents one mouse. Bars represent mean ± SEM

Whereas innate immune cells and T lymphocytes are well known to respond to IL-12 and IL-23 ^15,16^, the individual contribution of each cytokine to AD-driven neuroinflammation remains unresolved. Studies addressing the downstream effects of CNS-specific IL-12/23 signaling in the context of AD, as well as the unambiguous identification of their brain stromal targets are currently missing – a fact that is aggravated by the lack of reliable detection tools such as antibodies to most components of these multi-subunit cytokines as well as to their receptors. To systematically tackle these questions and dissect the precise mechanism of this yet unidentified neuroinflammatory pathway, we investigated the amyloid-carrying hippocampus in the presence or absence of IL-12/IL-23 (p40) signaling by means of an unbiased single-nucleus RNA sequencing (snRNA-seq) approach. We identified oligodendrocytes and distinct neuronal subtypes as the primary targets and downstream effectors cells of IL-12, while IL-23 did not seem to play a role in AD pathogenesis. Moreover, by intervening with IL-12 signaling, the AD-related loss of mature oligodendrocytes was rescued indicating that specific targeting the IL-12 signaling cascade is a promising interventional approach in the treatment of AD.

## Results

### Amyloid-β-dependent loss of oligodendrocytes is rescued upon deletion of IL-12/IL-23 signaling in the aged murine hippocampus

Given the central role of the hippocampus in contributing to cognitive functions and its involvement in AD-pathology^17^, characterized the transcriptional signature of individual hippocampi dissected from 250-day old amyloid-β-overexpressing AD-like mice harboring (APPPS1) or lacking IL-12/IL-23 signaling (APPPS1.*Il12b*^-/-^) as well as wildtype littermate control animals (WT) by single nucleus RNA sequencing. Three independent experiments (Fig. 1b) yielded a total of 82,298 nuclei expressing an average of 1,412 genes and 2,421 transcripts (defined as unique molecular identifiers (UMIs)), after removal of low-quality nuclei and doublets (Extended Data 1a-e). Roughly 86 % of all captured transcripts were protein coding while 13 % were comprised of long noncoding RNAs (lncRNAs), many of which were cell type-specific (Extended Data 1 f,g).

Unsupervised clustering followed by UMAP for visualization revealed 37 clusters which were assigned to various neuronal, glial and other cell types based on the expression of known marker genes (Fig. 1c, Extended Data 1h)^18,19^. We identified 18 clusters of excitatory neurons which were assigned to the anatomical region of the dentate gyrus, the cornu ammonis (CA) 1, CA2/CA3 and subiculum, 3 clusters of inhibitory neurons and 7 clusters of glial cells (microglia, astrocyte, oligodendrocytes, oligodendrocyte precursor cells (OPCs)), and a small cluster with features of Cajal-Retzius cells. Non-neural cells comprised myeloid cells such as peripheral macrophages (distinct from CNS-resident microglia) and small clusters of fibroblasts, choroid plexus and endothelial cells. The snRNA-seq data from all three independent biological replicates were superimposable, i.e. showed no major batch effects and justified data aggregation without resorting to batch correction or alignment procedures (Extended Data 2a-d). Furthermore, gene expression levels correlated between hippocampal snRNA- and bulk RNA-seq data (R ≥ 0.75) indicating that the single-nuclei data reflects the transcript composition of the intact tissue (Extended Data 3a). As expected, the *Il12b gene* was strongly deregulated in bulk RNA-seq data in APPPS1 compared to WT animals (Extended Data 3c). In total, when comparing bulk gene signatures of APPPS1.*Il12b*^-/-^versus APPPS1 mice, only five genes were shown to be significantly altered, i.e. either upregulated (*Pttg1*, *Gm15169*) or downregulated (*Il12b*, *Gm12663 and Gm24270*) (Extended Data 3d), highlighting the need of using an in-depth single-cell approach to study disease-related gene signatures.

The hippocampus as a particularly neuron-rich brain region revealed a majority of neurons amongst the recovered nuclei (60 % combined of excitatory and inhibitory neurons), followed by glia (37 % combining microglia, oligodendrocytes, astrocytes and OPCs (Extended Data 1i). These cell type proportions are consistent with the cell type composition of mouse hippocampus, e.g. as determined by the Blue Brain Cell Atlas (68 % neuronal versus 32 % of glial cells) (Extended Data 1j)^20,21^. A major pathological hallmark of AD is a substantial phenotypic alteration and proliferation of CNS-resident microglia. This was reflected in a higher number of microglial nuclei detected in the hippocampus of APPPS1 mice in comparison to their WT counterparts, irrespective of whether APPPS1 mice expressed or lacked IL-12/IL-23 signaling (Fig. 1d). Interestingly, we observed a substantial reduction of oligodendrocytes, but not of their precursors, in APPPS1 mice compared to WT mice. Notably, the AD-induced loss of oligodendrocytes was rescued in APPPS1.*Il12b*^-/-^ mice. Cell type deconvolution of bulk RNAseq data^22^ and quantification of oligodendrocytes in brain tissue sections by means of immunohistochemistry confirmed these findings (Fig. 1e-f). Olig2-positive cells exhibiting oligodendrocyte morphology were significantly reduced in APPPS1 mice, and this reduction was rescued to WT level in APPPS1.*Il12b*^-/-^ mice (Fig. 1g; Extended Data 3e-f)).

On the basis of a combination of the most highly variable genes in the oligodendrocyte lineage and all expressed transcription factors, we observed a differential trajectory of oligodendrocyte clusters resembling various stages of differentiation or maturation (Fig. 2a-b). A detailed analysis revealed that more mature oligodendrocytes, namely myelin forming oligodendrocytes (MFOL) and mature oligodendrocytes (MOL), accounted for the strong reduction of this cell population in APPPS1 mice (Fig. 2c-d). To assess whether the reduced number of mature oligodendrocytes might be the result of a dysregulation in OPC maturation in APPPS1 mice, we investigated their genetic profile based on differential gene expression (Extended Data 4a-c). Pseudotime analyses of genes involved in regulating oligodendrocyte differentiation positively or negatively revealed no gross difference between APPPS1 and APPPS1.*Il12b*^-/-^ mice across all oligodendrocyte maturation states (Extended Data 4d-g).

**Figure 2:**
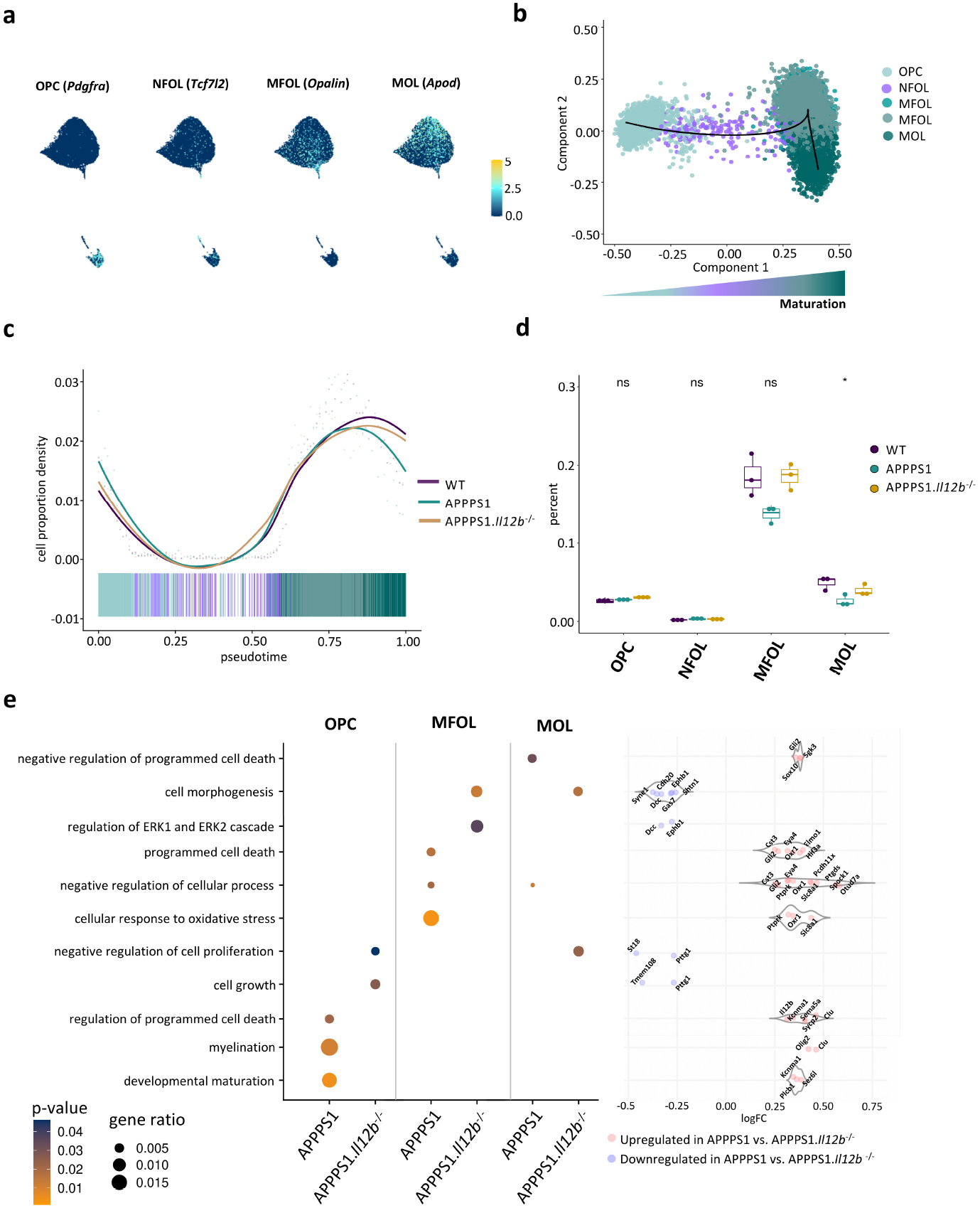
Mature Oligodendrocytes in the diseased APPPS1 hippocampus are susceptible to amyloid-driven neuroinflammation. **a)** Feature plots highlighting markers that characterize known oligodendrocyte maturation states. **b)** Pseudotemporal ordering of oligendrocytes revealed increasing differentiation along the known maturation trajectory from OPC via NFOL to MFOL and MOL. **c)** Cell proportion density along the pseudotime suggests an increasing loss of more mature oligodendrocytes in the diseased APPPS1 hippocampus. **d)** Loss of oligodendrocytes reaches statistical significance at the stage of MOL and is rescued by the absence of IL-12. **e)** Gene Ontology of differential expression between genotypes APPPS1 vs. APPPS1.*il12b*^-/-^. The dot size shows gene ratio and the color denotes p-value. Violin plot shows log2 fold changes of genes to corresponding GO term. Red dots indicate genes with positive log2 fold changes, blue dots indicate gene with negative log2 fold changes.

Gene ontology (GO) analysis in OPCs revealed enriched genes involved in myelin maturation (*Clu*, *Olig2*) and developmental differentiation (*Plcb1*, *Kcnma1*, *Sez6l*) in APPPS1 versus APPPS1.*Il12b*^-/-^ mice, indicating compensatory processes in OPCs directed at replacing dysfunctional and/or lost mature oligodendrocytes. In MFOLs of APPPS1 mice, genes involved in cellular responses to oxidative stress (*Slc8a1*, *Ptprk*, *Oxr1*) and programmed cell death (*Elmo1*, *Oxr1*, *Hif3a*, *Gli2*, *Eya4*, *Cst3*) were upregulated compared to MFOLs derived from APPPS1.*Il12b*^-/-^ mice, while three of those genes, namely *Gli2*, *Oxr1* and *Eya4*, also have been shown to inhibit cell death^23–25^. Similarly, APPPS1 MOLs showed an unexpected upregulation of genes inhibiting programmed cell death (*Sox10*, *Gli2*, *Sgk3*) compared to MOLs derived from APPPS1.*Il12b*^-/-^ mice (Fig. 2e), indicating that IL-12 signaling is capable of regulating oligodendrocyte homeostasis by mediating the balance between cell death and survival.

In summary, we show that IL-12/IL-23 signaling in Aβ-rich hippocampi of APPPS1 mice induces a loss of mature oligodendrocytes without affecting the number of OPCs and of other non-neuronal cell types such as microglia and astrocytes. Moreover, the decrease in oligodendrocytes within the amyloidogenic, inflamed microenvironment of the APPPS1 brain is driven by an IL-12 dependent cell death of MOLs rather than a global change in the maturation trajectory from the OPC population to fully differentiated oligodendrocytes.

### Neurons and oligodendrocytes act as IL-12 target cells in the Aβ-rich brain

In order to unravel how the thus far insufficiently understood IL-12/IL-23 pathway operates in the amyloidogenic brain, we determined the cell type-specific expression of IL-12/IL-23-associated transcripts within our snRNA-seq data. *Il12b* transcripts encoding for the common p40 subunit of IL-12/IL-23 signaling were induced and readily detectable in APPPS1 mice, but undetectable in WT and APPPS1.*Il12b*^-/-^ mice (Extended Data 5a). As expected from our earlier work, we found *Il12b* expression in the Aβ-harboring AD brain in microglia, but – somewhat unexpectedly – also in all other brain cell types, although at substantially lower levels (Fig. 3a). Even though *Il12b* was only expressed in the AD context, *Il12a* encoding the IL-12 subunit p35 (Extended Data 5b) was equally expressed across all three mouse genotypes, i.e. independent of AD pathology and irrespective of the presence or the lack of p40 signaling. *Il12a* transcripts were exclusively found in neurons and oligodendrocytes but – to our surprise – not in microglia (Extended Fig. 5c,d). Similarly, also the receptor subtypes were expressed at equal levels across all genotypes; *Il12rb1* was mostly expressed in oligodendrocytes (Fig. 3b), whereas *il12rb2* was more generally distributed across several cell types though more pronounced in neurons and oligodendrocytes (Fig. 3c). These findings are in line with single-nucleus sequencing data from Habib and colleagues describing the receptor subunit expression in neurons and oligodendrocytes^26^ (Extended Data 5e). Transcripts encoding the IL-23 receptor (*Il23r*) (Fig. 3d) and *Il23a* were barely detectable in the hippocampus (Extended Data 5f,g) and completely absent in oligodendrocytes, suggesting that IL-23 is not involved in mediating the p40-dependent changes described in APPPS1 mice^13^.

**Figure 3:**
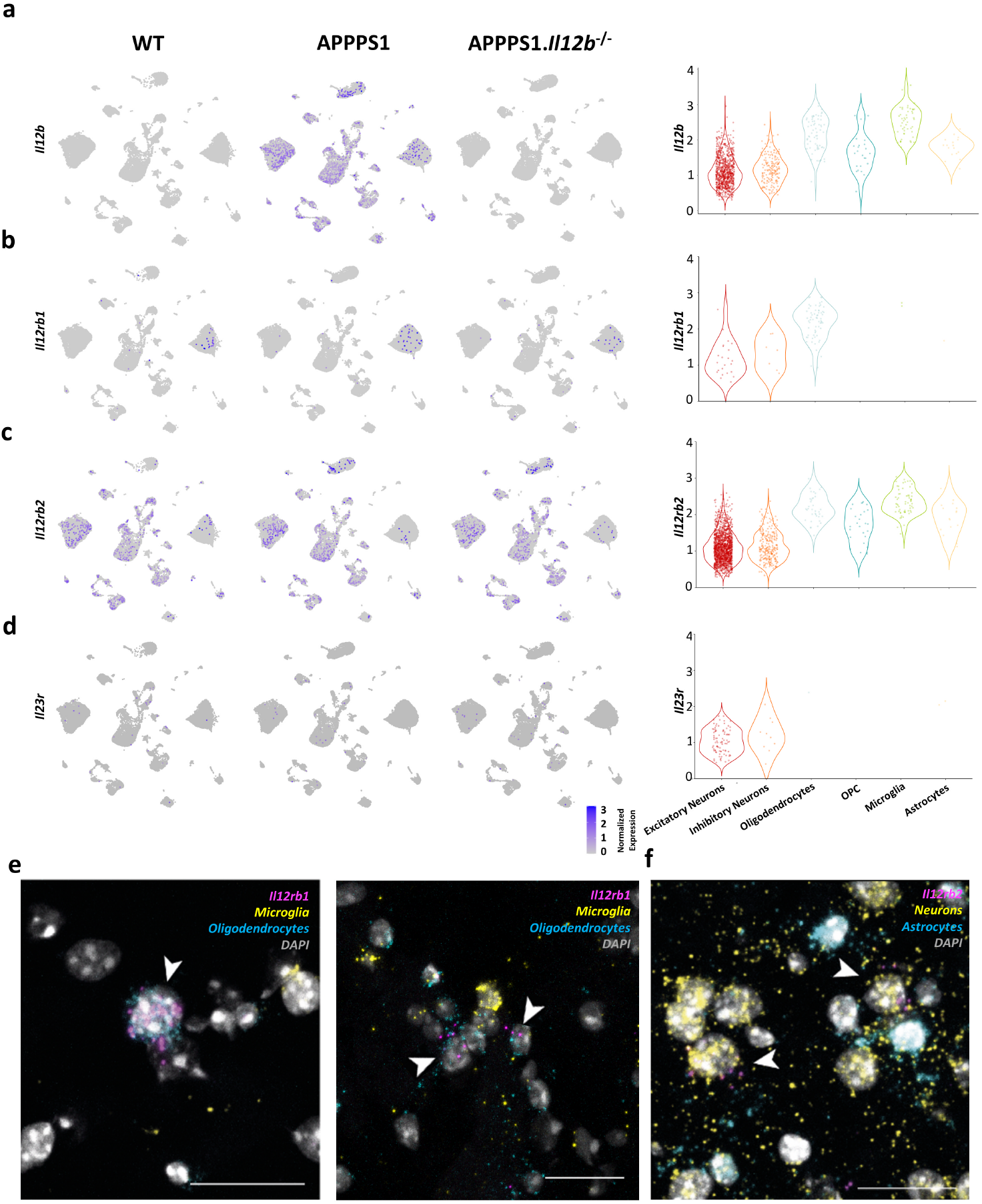
Oligodendrocytes and neurons express IL-12 receptor transcripts. **a)** *Il12b* expression. Featureplot showing *Il12b* is expressed disease specific and across all cell types. Violinplot depicting number of *Il12b* captured transcripts per cell type with zero counts removed **b)** *Ill2rbl*, coding for Interleukin-12 receptor subunit β1, is expressed across all three genotypes equally and mostly in oligodendrocytes. Violinplot showing captured *Il12rb1* transcripts **c)** *Il12rb2*, coding for Interleukin-12 receptor subunit β2, is strongly expressed in neurons, and to a lesser extent in microglia and oligodendrocytes. Violinplot showing captured *Il12rb2* transcripts. **d)** *Il23r* transcripts were almost absent in the aged mouse hippocampus. Violinplot showing captured *Il23r* transcripts. **i)** Combining probes for specific cell types with probes detecting the receptor subtypes, representative image of Single-molecule RNA fluorescence in situ hybridization in APPPS1 brain tissue revealed *Ill2rbl mRNA*-positive puncta (pink) in oligodendrocytes (marked in blue by expression of *sox10* mRNAs) and **f)** *Il12rb2 mRNA-*positive puncta in neurons (marked in yellow by *tub2/rbfox3 mRNAs*). *Il23r* puncta were not detectable. Scale bar= 25μm.

To visualize transcript expression, we applied single molecule RNA fluorescence *in situ* hybridization on aged mouse brain tissue, where each fluorescent spot corresponds to one single RNA transcript. This way, we were able to detect *Il12rb1*-positive RNA molecules localizing to *Sox10*-positive oligodendrocytes (Fig. 3e). *Il12rb2* RNA transcripts, however, were found within *Rbfox3/Tubb3*-positive neurons (Fig. 3f), while *Il23r* signals were not detectable, overall in line with the results retrieved by snRNASeq. Slight differences in detecting *Il12rb2* and *Il12rb1* transcripts e.g. in cells other than oligodendrocytes and/or neurons were due to the somewhat reduced sensitivity of the *in situ* hybridization compared to snRNASeq.

Taken together, our data illustrate that *Il12b* transcripts are predominately but not exclusively expressed by CNS-resident microglia in an amyloid-β-rich CNS microenvironment, while mainly oligodendrocytes, due to their expression of heterodimeric IL-12Rβ1/β2, harbor the molecular repertoire required to respond to increased IL-12/IL-23 levels in the pathological AD setting.

### Unaltered disease-associated microglia (DAM) signature in the absence of *Il12b* expression in AD-like brains

Since microglia are attributed to have a strong impact on neuroinflammation and AD pathology, we assessed transcriptional changes in microglia from APPPS1 mice with and without genetic ablation of *Il12b*. We observed two microglia clusters (Fig. 4a), one of which was prominent in APPPS1 and APPPS1.*Il12b*^-/-^ mice, i.e. AD specific, but virtually absent in age-matched WT control mouse brains. Differential gene expression analyses between these two microglia clusters showed upregulation of few *Trem2*-independent (*Apoe*) and many *Trem2*-dependent genes (*Ank*, *Csf1*, *Clec7a*, *Axl*, *Spp1*, *Itgax*, *Igf1*)^27^ including lipid metabolism and phagocytic pathways (*Lpl*, *Cst7*, and *Cd9*) as well as the downregulation of homeostatic microglia genes (*P2ry12*, *Cx3cr1* and *Tmem119*) (Fig. 4b). This gene expression signature showed strong similarities to the previously described signature from disease-associated microglia (DAM)^27^, which is linked to an altered microglial activation state in AD, including a modified phagocytotic capacity and secretion of neuroinflammatory mediators. Apart from an overlap with many previously described DAM genes (Extended Data 6a, Supplementary Table 1), we also identified significantly upregulated (*Ctnna3*, *Fgf13*, *Cacna1a*, *Olfr11*, *Ptchd1*, *Xylt1*) and downregulated genes (*Bank1*, *Nav2*, *Stab1*, *Zfhx3*, *Agmo, Ccr5*, *Elmo1*) in microglia of APPPS1 mice, which were not previously described as part of the DAM signature (Extended Data 6b, Supplementary Table 2).

**Figure 4.**
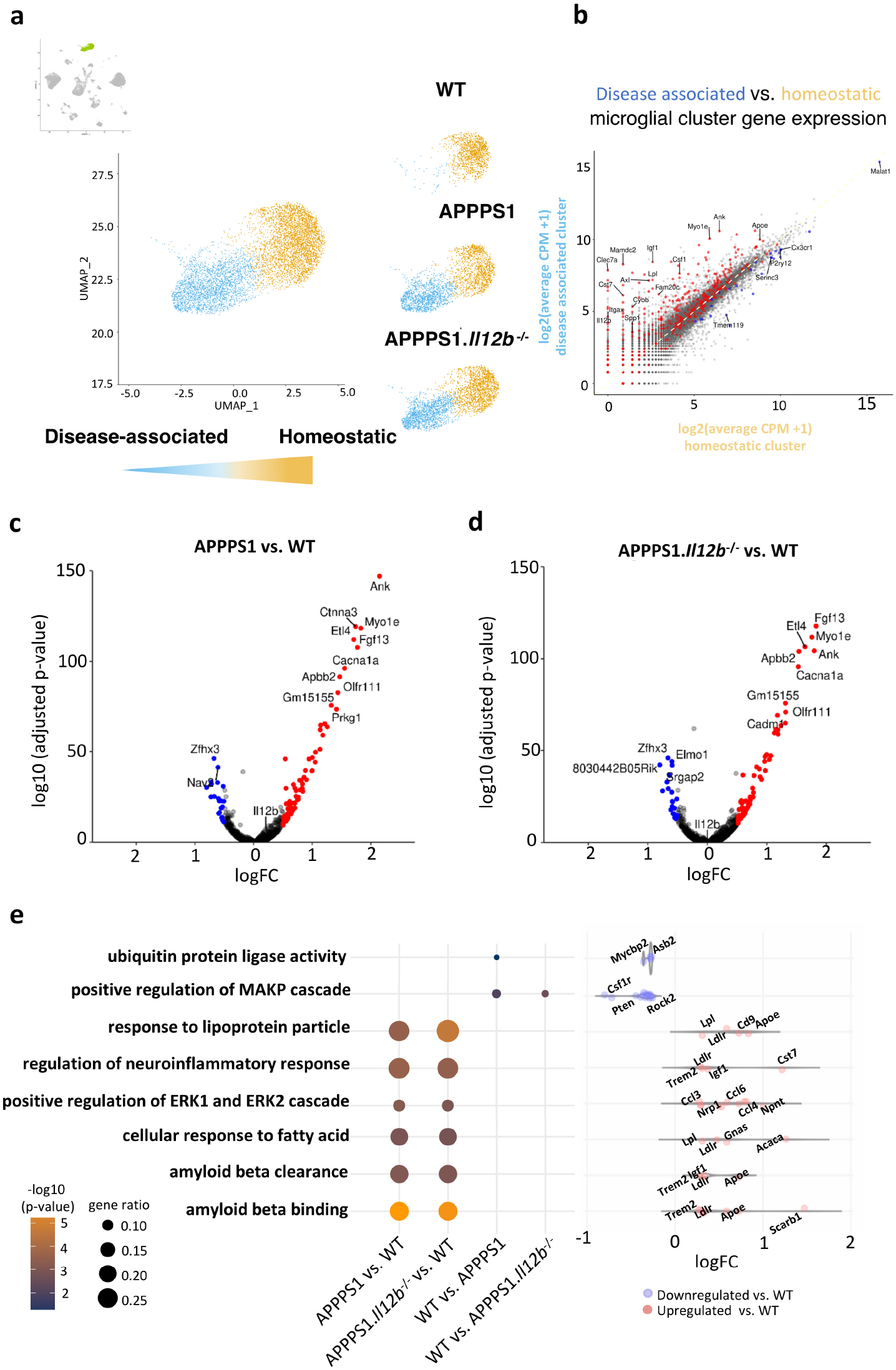
Microglia in APPPS1 and APPPS1.*Il12b*^-/-^ mice show gene signatures associated with enhanced microgliosis. **a)** Distinct homeostatic (yellow) and disease-associated (blue) microglia clusters were found in the combined snRNA-seq data set with the disease-associated clusters present only in APPPS1 and APPPS1.*Il12b*^-/-^ mice. **b)** Scatterplot comparing the gene expression in the disease-associated clusters versus the homeostatic clusters. **c)-d)** Volcanoplots comparing gene expression from all microglia in **c)** APPPS1 versus WT and **d)** APPPS1.*Il12b*^-/-^ vs WT. **e)** Gene Ontology analysis of differentially upregulated genes in the indicated genotypes. The dot size shows gene ratio and the color denotes p-value. Violin plot showing log2-fold change of certain specific genes to the corresponding GO term. Red dots indicate positive, blue dots negative log2-fold changes in APPPS1 and APPPS1.*Il12b*^-/-^ vs. WT.

Surprisingly, the inflammatory microglial gene signature of APPPS1 mice was largely unaffected in APPPS1 mice lacking IL-12/IL-23 signaling (Fig. 4c,d) resulting in a similar differential gene regulation of microglia from both APPPS1 and APPPS1.*Il12b*^-/-^ mice when compared to WT mice (Fig. 4e and Extended Data 6c). Only genes involved in ubiquitin protein ligase activity (*Mycbp2*, *Asb2*, *Rnf216*, *Rnf130*, *Znrf1*), thought to be linked to regulating neuroinflammation in AD^28^, were found to be slightly upregulated in WT vs. APPPS1, but not significantly in APPPS1.*Il12b*^-/-^ mice. However, it remains to be resolved whether these changes are of biological relevance, as we did not detect alterations in the neuroinflammatory profile in APPPS1 vs. APPPS1.*Il12b*^-/-^ mice.

Besides protein-coding RNAs, non-coding RNAs are part of the microglial immune response ^29,30^. Since the non-coding linear *Pvt1* and *circPvt1* as well as *Neat1* are capable of regulating the immune response ^31,32^, we assessed their expression in APPPS1 mice with or without functional p40 signaling. *Pvt1* and *Neat1* were equally and indistinguishably upregulated in APPPS1 and APPPS1.*Il12b*^-/-^ mice when compared to WT mice (Extended Data 6d-h), corroborating our finding that the disease-associated coding and noncoding gene signature of microglia in amyloid-β bearing APPPS1 mice is independent of IL-12 in the hippocampus.

### IL-12 signaling affects neuronal homeostasis preferentially in the subiculum of AD-like mice

When investigating whether IL-12 signaling affects neurons, we found the reduction in subicular neurons in AD-like APPPS1 mice compared to WT littermates (which did not yield significance) not altered by the lack of *il2b* (Extended Data 7a). However, of note were alterations in gene expression in subicular neurons of APPPS1 mice lacking *il12b* compared to those with functional IL-12 signaling (Extended Data 7b) known to be involved in pathways impacting hippocampal memory and synaptic plasticity such as *Erbb4 and Rarb*^33,34^. Besides seeing some differential regulation of genes involved in ion homeostasis, we also found an upregulation of genes implicated in dendrite development (*Dab1*, *Fat3*, *Fezf2*, *Fmn1*, *Hecw2*, *Klhl1*, *Map1b*, *Nedd3l*, *Sez6*, *Ss181*) in APPPS1.*Il12b*^-/-^mice, suggesting IL-12-dependent enhanced compensatory efforts aimed at regenerating neuronal homeostasis in the AD microenvironment (Extended Data 7c).

To gain insights into cell-cell communication between different neuronal subtypes, we applied cellphoneDB, a publicly available repository of distinct receptors, ligands and their interactions, to infer co-expressed receptor-ligand pairs derived from single-cell RNA sequencing data^35,36^. This way we observed an IL-12-dependent alteration in the receptor ligand pairing of neuropilin (*Nrp*)1, *Nrp2* and vascular endothelial growth factor A (*Vegfa*) as well as *Ephb2* and *Grin2b* in excitatory neurons and inhibitory neurons in APPPS1 mice (Extended Data 7d) region-specifically including subiculum, dentate gyrus, and CA2/CA3. Nrp1 and Nrp2 serve as co-receptors for VEGF receptors and support neuronal guidance as well as dendritic growth and branching in the adult brain ^37,38^. Given the reduced Nrp1/2 signaling in APPPS1 vs WT and APPPS1.*Il12b*^-/-^ mice, the reduced synaptic density known to occur in aged APPPS1 mice may explain some of the cognitive alterations in these AD-like mice, which appear to be rescued by interfering with IL-12 signaling^13^. Another mechanism affecting neuronal homeostasis is based on alterations in Neuregulin (Nrg)-ErbbB signaling, which links oligodendrocyte and neuronal interaction and mediates a plethora of cellular functions in both cell types^39^, including Nrg1-ErbB-driven regulation of axonal pathfinding, synaptic behavior and neuronal migration^40,41^. Of note, CellphoneDB-based analyses revealed a reduced *Nrg1-Erbb3* interaction in newly formed oligodendrocytes (NFOL) in the dentate gyrus, CA1 and in the subiculum of APPPS1 compared to WT mice, which was rescued in APPPS1.*Il12b*^-/-^ mice, reaching WT levels (Extended Fig. 7e). This signaling affects oligodendrocyte survival *in vitro^42^* and has been shown to be key for proper myelination^43^. Similarly, Nrg2-Erbb4 was found to be reduced in dentate gyrus-derived MOLs of APPPS1 mice, which was reverted to WT levels in APPPS1.*Il12b*^-/-^ mice lacking functional *il12b* signaling. These data indicate that IL-12 signaling-dependent perturbations in the transcriptional profile of neurons (ultimately leading to functional dysregulation) might be mediated either through binding of IL-12 to its receptor directly on the neuronal cell surface, or through affecting oligodendrocytes IL-12-specifically leading to an alteration in their trophic support of neurons, or a combination thereof.

## Discussion

Given that interfering with IL-12/IL-23 signaling potently reverts AD pathology^13,14^, we aimed at precisely dissecting the molecular underpinnings of this immune cascade by using a well-established amyloidogenic AD-like mouse model^44^. Based on unbiased single-nucleus RNA sequencing followed by distinct validation approaches we can show that IL-12/IL-23 signaling in the AD-like CNS is executed exclusively through IL-12, and not through IL-23. IL-12 actions preferentially target oligodendrocytes and neurons, which harbor the respective transcripts for IL-12 receptors. IL-12 receptor subtypes were equally expressed in AD-like and WT mice, while only *Il12b* encoding for IL-12 ligand expression appeared to be AD-specific.

While earlier studies suggested a disease-related microglia-specific expression of IL-12/IL-23 (p40), our data – owing to the technical advantages of applying snRNA-seq - reveals that there is an AD-driven expression of IL-12 also in other CNS-intrinsic cells at later stages of AD pathology. These findings suggest that IL-12 is part of a damage-associated response in a variety of brain cells including microglia, astrocytes, oligodendrocytes and neurons. However, *Il12a*, coding for p35, the binding partner of p40, was not detectable in microglia, but was found to be expressed in oligodendrocytes and neurons at equal levels across all genotypes investigated, i.e. in AD-like mice with or without functional IL-12/IL-23 signaling and in WT mice. Only co-expression of both p35 (*Il12a*) and p40 (*Il12b*) results in the formation of bioactive IL-12p70; in the absence of p40, p35 misfolds and degrades^45,46^. Thus, only oligodendrocytes and neurons, which express both *II12a* and *II12a*, are molecularly equipped to produce soluble, functional IL-12p70, while microglia appear to produce p40 homodimers (IL-12p80) *in vivo*.

Notably, we discovered an IL-12/IL-23-dependent loss of mature oligodendrocytes in Aβ-overexpressing APPPS1 mice. While there is general agreement on the occurrence of white matter changes and myelin pathology in human AD patients based on imaging and postmortem data^47^, animal studies using AD-like mice are inconclusive in this respect: whilst *in vivo* studies suggest that oligodendrocytes may be affected by Aβ burden resulting in a decrease in myelin basic protein (MBP)^48,49^, other studies report that Aβ exposure to oligodendrocytes *in vitro* induces a more mature phenotype as evidenced by an increase in oligodendrocyte branching and MBP production^50^. By comparing the overall oligodendrocyte numbers in AD-like mice and AD patients, it has been suggested that the change in numbers reflects their dynamic alterations depending on disease state^51^. Changes in the number of oligodendrocytes in an amyloidogenic environment have been attributed to an imbalance in OPC proliferation and maturation or their differentiation^52^. While previous studies reported an increase in Olig2-cells in the cortex of Aβ overexpressing APPPS1^51^ and 5xFAD mice^53^, we found a substantial reduction in oligodendrocytes in the hippocampus of amyloid-harboring and neuro-inflamed APPPS1 mice, which is entirely rescued and reverted to WT levels upon deleting IL-12. On a transcriptional level, we found no gross alterations in genes involved in regulating oligodendrocyte differentiation and maturation, while mature oligodendrocytes – presumably based on their IL-12/IL-23 signaling repertoire - were affected by the AD-specific amyloidogenic and inflammatory microenvironment. Given the limited knowledge on how Aβ affects oligodendrocytes, but also how oligodendrocyte homeostasis may impact amyloid burden and AD pathology itself, the mechanism by which oligodendrocytes get rescued through inhibition of IL-12 signaling in APPPS1 mice deserves further research.

Besides substantially affecting oligodendrocytes, IL-12/IL-23 signaling also altered the gene signatures of subicular neurons of the hippocampus – a brain region of high relevance for cognitive functions. Atrophy in the subiculum is thought to be the earliest sign of neuronal degeneration in AD^54^ and is connected to memory loss^55^. While subicular neurons seemed to be decreased in APPPS1 mice lacking *Il12b*, their transcriptional signature was altered for genes involved in memory and synaptic plasticity such as *Erbb4* and *Rarb*, as well as genes regulating dendrite development such as *Nrp1* and *Nrp2*. The latter encode Neuropilin-1 and −2, which are known to engage with VEGF receptors to support neuronal guidance as well as dendritic growth and branching. The previously reported IL-12/IL-23-mediated rescue of cognitive deficits of APPPS1 mice^13^ may thus be explained by specific interference of IL-12 signaling with neuronal plasticity. Given the pronounced oligodendrocyte phenotype in APPPS1 mice, however, it cannot be excluded that such neuronal gene alterations are a secondary, indirect effect disturbing the close functional relationship between neurons and oligodendrocytes, e.g. due to a lack in trophic support of neurons by oligodendrocytes. In this context it is noteworthy to mention that the *Nrg1-Erbb3* interaction known to link oligodendrocyte and neuron functions, i.e. by modulating synaptic plasticity^56^, was found to be decreased in APPPS1 mice, but was rescued to WT levels in APPPS1 mice lacking IL-12 signaling.

Another remarkable phenotype we observed is that the lack of IL-12/IL-23 signaling in AD-like APPPS1 mice did not change the inflammatory state of IL-12/IL-23(p40)-producing microglia, including the respective DAM signatures. Thrupp *et al*. reported that snRNA-seq cannot fully recapitulate the DAM signature in (human) microglia ^57^, however multiple studies showed strong correlation between mouse snRNA-seq and single cell RNA-seq data^53,58–60^. Inflammatory transcriptional DAM signatures derived from protein coding as well as non-coding RNAs are thought to cause many pathological changes in microglia, from displaying a more neuroinflammatory profile to exhibiting an altered phagocytic capacity – as observed in AD. The fact that deletion of IL-12/IL-23 did not alter the DAM signature can be explained by paracrine signaling of microglia, which affects other IL-12/IL-23 recipient-prone CNS-intrinsic cells but spares to alter their own intrinsic functions. This way, IL-12/IL-23 inhibition provides an attractive way to selectively interfere with AD pathogenesis by targeting detrimental IL-12/IL-23 functions in non-microglial CNS-intrinsic cells whilst not affecting the phenotype of microglia, thus avoiding the risk of “ locking” microglia in either their homeostatic or disease-associated state.

In conclusion, our data is not only instrumental in dissecting the mechanism of IL-12/IL-23-specific immunomodulation of AD and in identifying its cellular targets, namely oligodendrocytes and neurons, but also highlights the potential of an IL-12/IL-23 targeted immunotherapy in AD. The fact that IL-12, but not IL-23 is the pathogenetically relevant pathway in AD-related IL-12/IL-23 signaling may either encourage the use of exclusive IL-12 inhibition in tackling AD, or highlights the minimized potential side effects when applying combined IL-12/IL-23 blockers – questions that need to be addressed in future studies, which will also need to investigate whether this newly identified neuroimmune crosstalk between various CNS-intrinsic cell types may also play a role in CNS diseases other than AD.

## Methods

### Mice

Heterozygous APPPS1^+/-^ mice (previously described by Radde and colleagues^44^, termed APPPS1 mice throughout this manuscript) were crossed to *Il12b*^-/-^ ^61^ mice and were compared to wild type littermate controls. Mice were bred on a pure C57BL/6J background. Animals were kept in individually ventilated cages with a 12 h light cycle with food and water ad libitum. All animal experiments were conducted in accordance with animal welfare acts and were approved by the regional office for health and social service in Berlin (LaGeSo; license O 298/17 and T 0276/07).

### Nuclei preparation

Mouse hippocampi were harvested from male mice at the age of 250 days and immediately snap frozen in liquid nitrogen and stored at −80°C until further processed for nuclei isolation. Nuclei were isolated from a single mouse hippocampus in 2 ml of pre-chilled EZ PREP lysis buffer (NUC-101, Sigma) using a glass Dounce tissue grinder (D8938, Sigma) (25 strokes with pastel A and 25 strokes with pastel B) followed by incubation for 5 minutes on ice with additional 2 ml of EZ PREP buffer. During incubation, 1 μM DAPI was added to the homogenate. The homogenate was then filtered through a 30 μM FACS tube filter. A BD FACSAria III Flow Cytometer with a 70 μm nozzle configuration was used to sort the nuclei based on the fluorescent DAPI signal at 4°C. As CNS nuclei vary strongly in size, no doublet discrimination was performed based on FSC or SSC to avoid bias against nucleus size. Nuclei were then counted based on brightfield image and DAPI fluorescence using a Neubauer counting chamber and a Keyence BZX-710 microscope.

### Single-nucleus RNA sequencing

Isolated mouse nuclei were immediately used for droplet-based 3’end single-cell RNA sequencing using the Chromium Next GEM Single Cell 3’ GEM, Library & Gel Bead Kit v3.1, following the manufacturer’s instructions (PN-1000121, 10x Genomics). The libraries were multiplexed and three samples per lane were sequenced on an Illumina HiSeq 4000 sequencer. Demultiplexing, barcode processing, read alignment and gene expression quantification was carried out using Cell Ranger software (v3.1.0, 10x Genomics). First, cellranger mkfastq demultiplexed the sequencing by sample index. The quality of the data was checked using FastQC (v0.11.5) and all samples showed high quality RNAseq data with good median per-base quality (≥28) across most of the read length. Cellranger count used STAR software with default parameters to align sequenced reads to the reference genome (GRCm38, Ensembl GTF version 98). As nuclei have a high amount of pre-mRNA, we generated a custom pre-mRNA reference based with the instructions provided on the 10X Genomics website; we also included intronic reads in the final gene expression counts. Finally, the output files for all 9 samples were aggregated into one gene-cell matrix using cellranger aggr without read depth normalization.

### Quality control and data pre-processing

Data was analyzed in R (v3.6.0) using Seurat (v3.1.2)^62^. In all downstream analyses, the filtered feature-barcode matrix was used rather than the raw feature-barcode matrix. For the initial quality control, we excluded genes expressed in less than 3 nuclei and nuclei expressing less than 200 genes or less than 500 or more than 30,000 UMIs and nuclei with more than 5% mitochondrial reads. This resulted in a dataset of 84,002 cells and 31,790 quantified genes across 9 samples.

### Dimensionality reduction, clustering, visualization and cell type identification

After initial quality control, we normalized UMI counts utilizing the “ LogNormalize” method and by applying a scale factor of 10,000. We found variable genes using “ FindVariableFeatures” with the selection method “ vst”. In addition, data regression was performed using the ScaleData function with nUMI, percent mitochondrial counts, and sample origin as confounding factors. Dimensionality reduction was performed using PCA and we selected 40 PCs based on Elbow plot. The FindClusters function, which implements shared nearest neighbor (SNN) modularity optimization-based clustering algorithm was applied with a resolution of 0.8 and identified in 45 initial clusters. A further dimensionality reduction step was carried out, using the UMAP algorithm to visualize the data. The UMAP parameters were: n.neighbors = 20, min.dist = 0.35, n.epochs = 500, spread = 2. As UMAP overlay by individual sample shows minimal batch effects, we didn’t consider any batch correction method. For assigning clusters to cell types, we used, the “ FindAllMarkers” function with default parameters was used, identifying negative and positive markers for that cluster.

### Cell doublet identification

Scrublet (v0.21)^63^ with an expected_doublet_rate = 0.06 was applied, resulting in detection of 5.2% of doublets. We defined doublet clusters as containing more than 50% of doublets and removed these for downstream analysis. We noticed that these clusters were projected in the middle of other cell types in UMAP and could be validated as expressing marker genes from two different cell types. In the end, we removed 8 clusters, which reduced our data set to 82,298 nuclei.

### Relative entropy

Cell type variability was measured using an entropy-based approach^64^. We first grouped by replicate and genotype. The local neighborhood was defined by taking the 30 nearest neighbors using kNN-Graph and the relative entropy. We applied Kullback-Leibler divergence to measure how cells are distributed among samples. Controls were randomly shuffled and showed that differences detected in gene signature were of biological relevance and not driven by technical artifacts.

### Cellular proportions

One-way ANOVA was used to test whether cellular proportions differed by genotype. Homogeneity of variance and normality of data distribution was assessed by using Bartlett and Shapiro-Wilk tests, respectively, with the R package stats (v3.6.0). A p-value of less than 0.05 was considered statistically significant.

### Identification of differentially regulated gene expression

In order to identify differentially regulated genes among genotypes, we performed the empirical Bayes quasi-likelihood F-tests (QLF) including the cellular detection rate (the fraction of detected genes per cell) utilizing EdgeR (v3.28.1)^65,66^. A log2-fold change greater than 0.25 and an FDR less than 0.01 were considered significant. Among differentially expressed genes, we removed the *Ttr* gene as its expression was highly dependent on the presence of a Choroid Plexus cluster in a given sample, suggesting a dissection bias at the stage of hippocampus isolation.

### Gene ontology (GO) analysis

GO term enrichment of each cluster was performed with topGO (v2.36.0)^67^. In GO analysis, genes showing average log2 fold change > 0.25 and adjusted p value < 0.01 were considered significant and all expressed genes were used as background. We used the elim algorithm instead of the classic method to be more conservative and excluded broad GO terms with more than 1000 listed reference genes.

### Trajectory inference analysis

We performed trajectory inference with SCORPIUS (v1.0.7)^68^ for oligodendrocytes populations including OPCs. To infer developmental trajectory, we used highly variable and all expressed transcriptional factors genes and reduced dimension using distance metric as spearman with 3 number of dimensions. To infer gene expression along the pseudotime, we first downloaded a list of genes from Mouse Genome Informatics (http://www.informatics.jax.org/) displaying negative regulation of oligodendrocyte differentiation (GO:0048715) and positive regulation of oligodendrocyte differentiation (GO:0048714), which was smoothed over pseudotime using generalized additive model using mgcv (v1.8-28).

### Cell-cell Interaction

We used CellPhoneDB (V2.1.1)^35,36^ to assess cellular crosstalk between different cell types. To identify putative cell-cell interactions via multi-subunit ligand-receptor complex pairs, label permutation was performed. First, we converted mouse gene symbols to human gene symbols using biomaRt (v2.42.1)^69^ and removed duplicated gene symbols from digital gene expression matrix. We then calculated normalized data with scale factor 10,000. Finally, we conducted statistical analyses by randomly permuting the cluster labels of each cell 10,000 times.

### Comparison with genes expressed in disease-associated microglia (DAM)

A list of DAM genes was downloaded from supplementary table 2 of the work by Keren-Shaul and colleagues^27^. This DAM signature were collected from single-cell sorting and downstream single-cell RNA-seq of microglia from the AD-mouse model 5xFAD. We then computed the log2 fold changes of the Microglia 3 (Disease-associated cluster) to Microglia 1 (Homeostatic cluster) ratio for each gene^27^, resulting in 461 DAM genes. To identify DAM APPPS1-related signature genes from our dataset, we compared the differential expression of cluster 8 (disease state) and cluster 3 (homeostatic state), where only logFC > 0.25 and FDR < 0.01 were considered, ultimately resulting in 488 genes. 96 genes intersected between our study and the already published study^27^. Of those, 365 genes were specific for the published DAM signature, while 392 genes were specific to our APPPS1-related dataset. While there seems to be an ubiquitous gene signature related to microglioses in AD-related murine mouse models, our study also highlights the specific microglial gene signature of APPPS1 mice.

### Visualization of read coverage with genomic signal tracks

To visualize read coverage of single-nucleus RNA sequencing data in a genome browser, Sambamba (v0.6.8)^70^ was used to sort BAM file produced from 10X cellranger count. We extracted only primary alignment reads from sorted BAM file and created bedgraph file using bedtools (v2.27.1)^71^ with normalized using read depth and split file by strand specific. Finally, we created a BigWig file using bedGraphToBigWig (v4)^72^ and the resulting genomic signal tracks were visualized in the UCSC Genome Browser. The results of genome tracks is located in public hubs at MDC Genome web browser (https://genome.mdc-berlin.de/)

### Bulk-RNA isolation

Total RNA was isolated from freshly frozen hippocampus from 250 day old animals using the NucleoSpin miRNA and RNA purification kit (740971.50, Marchery Nagel). In short, the tissue was homogenized using a Pellet Mixer (431-0100, VWR) in 0.35 ml Buffer ML provided in RNA purification Kit and subsequently passing the homogenate through a G23 needle (465 7667, B. Braun) until no clumps remained. RNA was isolated according to manufacturer’s protocol and eluted in 20 μl of RNAse-free water.

### Bulk transcriptome analysis

Library construction and bulk mRNA-seq was performed by Novogen (UK) Company Ltd. (non-stranded cDNA libraries; 150 bp paired-end run with a depth of 40 million reads per library). Bulk transcriptomes were aligned using STAR (v2.7.1a)^73^ with mm10 reference and quantified using featureCounts (v1.6.0). Differential expression genes between samples were determined as adjusted P value less than 0.05 and fold change higher than 1 or lower than −1 using DESeq2 (v1.24.0)^74^ without the lfcShrink function. To analyze pairwise correlations between bulk transcriptomes and single-nuclei RNA-seq data, bulk transcriptomes were converted to transcripts per million (TPM) by dividing each counts of each gene by its length and multiplying by one million. Each gene length was calculated from GTFtools(v0.6.9)^75^ by median length of its isoforms. Single-nuclei RNA-seq expression counts were summed by each sample and converted to counts per million (CPM). The scale of all figures as log2 (CPM/TPM + 1).

### Cell type deconvolution

To validate the findings provided by bulk transcriptomics regarding the ratio of various CNS cell types, we performed weighted non-negative least squares for cell type proportion estimation utilizing Multi-subject Single Cell deconvolution (MuSic (v0.1.1)^22^. We ran the package with default settings with highly variable genes from snRNA-seq data.

### Multiplex single-molecule RNA fluorescence in situ hybridization (FISH)

Frozen brain tissue was placed in a tissue mold (SA62534-15, Sakura) and submerged in Tissue-Tek freezing medium (4583, Sakura). 10 μm thick tissue sections were cut using a cryostat (Thermo Scientific HM 560), placed on SuperFrost Plus slides (500621, R. Langenbrink) and dried for 1 hour at −20°C. Tissue processing for RNAscope^®^ multiplex staining (Advanced Cell Diagnostics, Inc.) was done following manufacturer’s protocol for fresh frozen sections. In brief, tissue was fixed in freshly prepared 4 % PFA (pH 7.4) for 30 minutes at 4°C, followed by alcohol dehydration. Tissue was exposed at room temperature to the given concentration of H_2_O_2_ for 10 minutes and to Protease IV (322340, Bio-Techne) for 30 minutes and then incubated for two hours with target probes at 40 ° C in a HybEZ^™^ Hybridisation System (321711, Bio-Techne). The following target probes were used: Mm-Il12rb1 (488761, Bio-Techne), Mm-Il12rb2 (451301, Bio-Techne), Mm-Il23r (403751, Bio-Techne), Mm-Aldh1l1-C2 (405891-C2, Bio-Techne), Mm-Slc1a3-C2 (430781-C2, Bio-Techne), Mm-Gfap-C2 (313211-C2, Bio-Techne), Mm-Sox10-C2 (435931-C2, Bio-Techne), Mm-Tmem119-C3 (472901-C3, Bio-Techne), Mm-Sall1-C3 (469661-C3, Bio-Techne), Mm-Rbfox3-C3 (313311-C3, Bio-Techne), Mm-Map2-C3 (431151-C3, Bio-Techne). Signal amplification was achieved using the RNAscope^®^ Multiplex Fluorescent Kit v2 (323110, Bio-Techne), closely following the manufacturer’s protocol. Probes were labelled with Opal^™^ 520 (1:500, C2 probe, FP1487001KT, Perkin Elmer), Opal^™^ 570 (1:500, C1 probe, FP1488001KT, Perkin Elmer) and Opal^™^ 690 (1:500, C3 probe, FP1497001KT, Perkin Elmer) and three-dimensional image stacks (1 μm step size, 40x objective) of stained sections were taken on a Leica TCS SP5 confocal laser scanning microscope using a HCX PL APO lambda blue 63× oil UV objective controlled by LAS AF scan software (Leica Microsystems).

### Immunohistochemistry

Animals were perfused with PBS and hemispheres were fixed for 24 hours in 4 % PFA at 4°C. Brains were further processed by incubating them in 10 %, 20 % and finally 30 % sucrose in PBS over the course of three days. Free-floating brain sections were cut at 30 μM thickness using a cryostat (NX70 957030L, Thermo Fischer) and stored in cryoprotectant (0.65 g NaH_2_PO_4_ × H_2_O, 2.8 g Na_2_HPO_4_ in 250 ml ddH_2_O, pH 7.4 with 150 ml ethylene glycol, 125 ml glycerine) at 4°C until further use. Sections were stained by first incubating them in 0.3 % Triton-X in PBS with 10% normal goat serum for 1 hour. The primary antibodies used for detecting oligodendrocytes was Olig2 (Rabbit, 1:750, AB9610, Milipore) and incubated at 4°C overnight. The secondary antibody (Alexa568 goat anti rabbit, 1:300, A11011, Invitrogen) was added for 2 hours at room temperature. Nuclei were counterstained using 500nM DAPI for 1 minute.

### Quantification Olig2-positive cells in brain sections

3-6 brain sections per animal, stained with Olig2 antibodies and DAPI, were imaged using an inverted microscope (Leica DMI 6000) with 40x magnification. Images were automatically stitched together using LAS X 3.3 Stage Experiment Tilescan software (Leica). Using the DAPI stained image, the hippocampus was set as region of interest (ROI). Within this ROI, using ImageJ with a set threshold for all images across all animals, all DAPI as well as Olig2-positive puncta were quantified using the “ Analyze Particle” function. The total count of Olig2-positive puncta were normalized to the total counts of DAPI-positive puncta.

## Additional Resources

We created a shiny app which allows to access our single-cell data for any gene of interest interactively via the following URL: https://shiny.mdc-berlin.de/AD_Neuroinflammation/.

## Code availability

Custom R scripts used to analyze data and generate figures are available upon request.

## Non-author contributions

We are indebted to Eileen Benke, Alexander Haake (Department of Neuropathology, Charité - Universitätsmedizin Berlin, Germany) for excellent technical assistance and advice. Cartoon images were partially created with Biorender.com. We thank Markus Landthaler and Laleh Haghverdi (BIMSB, MDC, Berlin, Germany) for advice. We thank Cledi Cerda Jara, Agnzieska Rybak-Wolf, Miriam Wandres, Anna Löwa (all BIMSB, MDC, Berlin, Germany) for fruitful discussions.

## Author Contributions

S.S., P.E., A.B., and C.B. performed and C.K., F.L.H and N.R. supervised experiments, S.J.K. performed and N.R. and N.K. supervised bioinformatics analyses. All authors contributed to the experiments and supported data analyses; N.R. and F.L.H. designed the study and procured funding. C. K., N.R. and F.H. jointly supervised the study; S.J.K. and S.S. prepared figures. S.S., S.J.K., P.E., B.B., N.K., C.K., N.R., M.A. and F.L.H. wrote and revised the manuscript. All authors approved the manuscript.

## Data and materials availability

All data is available in the main text or the supplementary materials.

