## Extended Data for "The neuroinflammatory interleukin-12 signaling pathway drives Alzheimer’s disease-like pathology by perturbing oligodendrocyte survival and neuronal homeostasis"

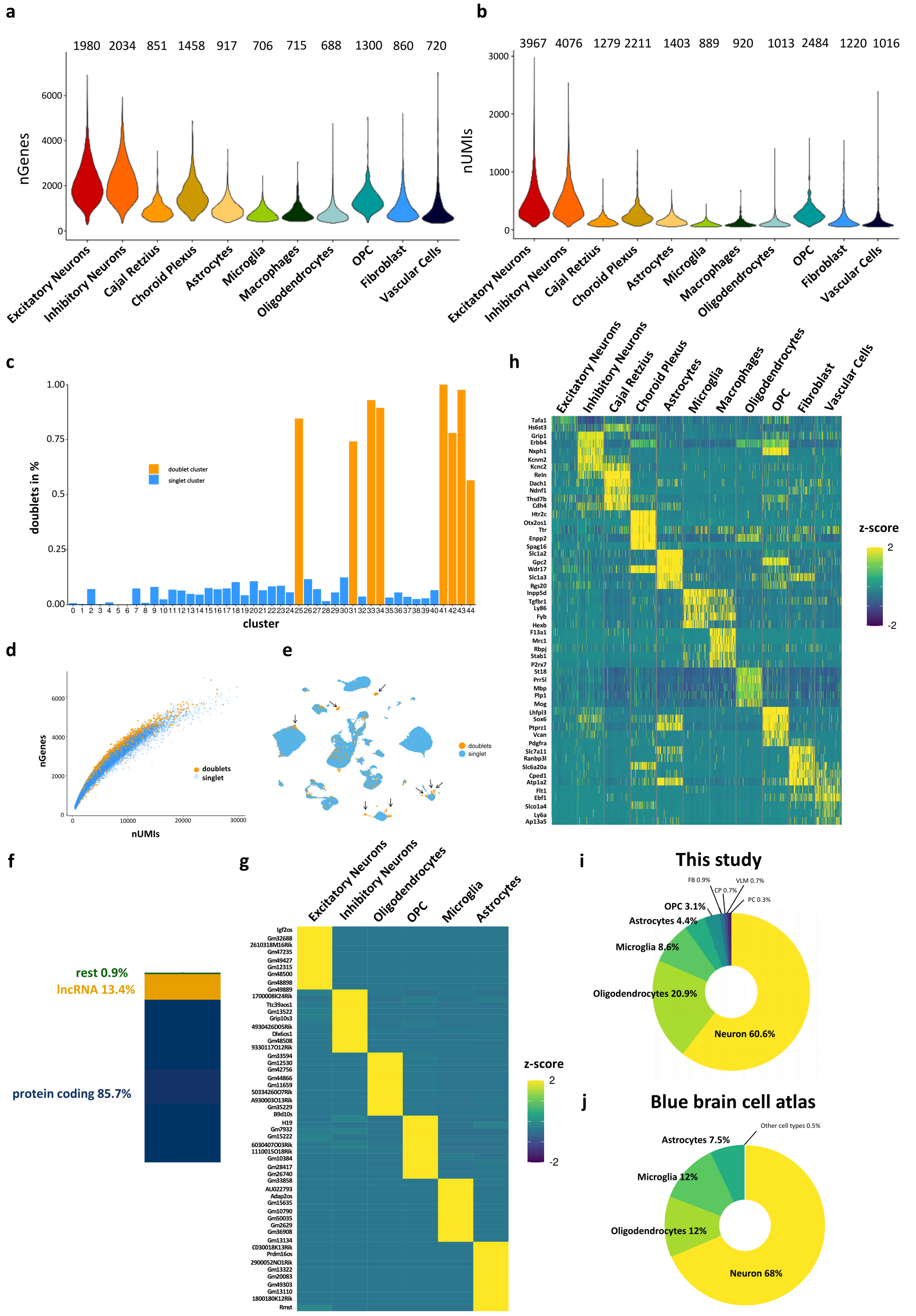


Figure legend on following page

**Extended Data 1: Data quality and cluster annotation. a),b)** Violin plot showing for different hippocampal cell types **a)** gene count per cell and **b)** UMI count per cell, averaging at 1,412 genes and 2,421 UMI, respectively. **c)-e)** Doublet discrimination identified roughly 5% of all nuclei as doublets. From a total of 44 detected clusters, cluster 25, 31, 33, 34, 41, 42, 42 and 44 were flagged as clusters carrying doublets using Scrublet (v0.21)[63]. Clusters with >50% of doublets were removed from downstream analysis. **f)** Bar graph depicting biotypes of detected RNA shows that approximately 85.7% of all transcripts were protein coding followed by 13.4% long non-coding RNAs (lncRNAs) **g)** Heatmap showing celltype specific lncRNAs **h)** Known celltype-specific gene signatures were used to annotate clusters[18, 19] **i)** Cellular proportions of this snRNAseq study reflected the **i)** published cellular composition of the hippocampus according to the Blue Brain Cell Atlas[20, 21].

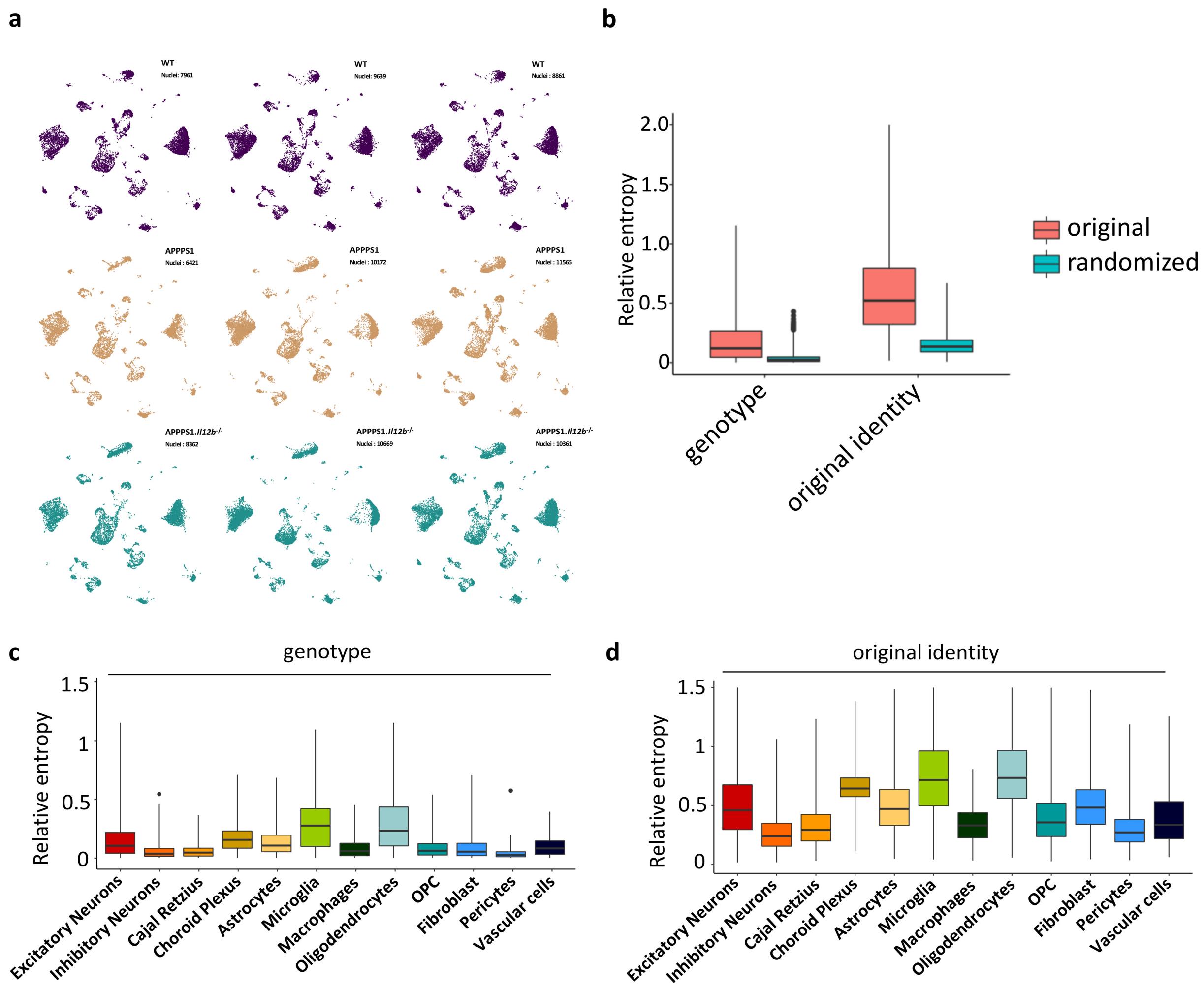


**Extended Data 2: Absence of major batch effects in hippocampal snRNAseq data from 250 day old mice. a)** UMAP visualization of three individual biological replicates per genotype, showing mostly equal representations in each cluster. **b) – d)** Entropy-based quantification of batch effects. **b)** The distribution of relative entropy values for cells grouped by sample (original identity) or genotype was greater than randomizing these labels across cells. **c)-d)** Different cell types grouped by **c)** genotype and **d)** sample (original identity). Relative entropy per cell type was highest for cell types that differed biologically between genotypes, for example microglia and oligodendrocytes which react most strongly to amyloid and inflammatory conditions. Thus, differences were driven by biological meaningful distinctions rather than technical errors.

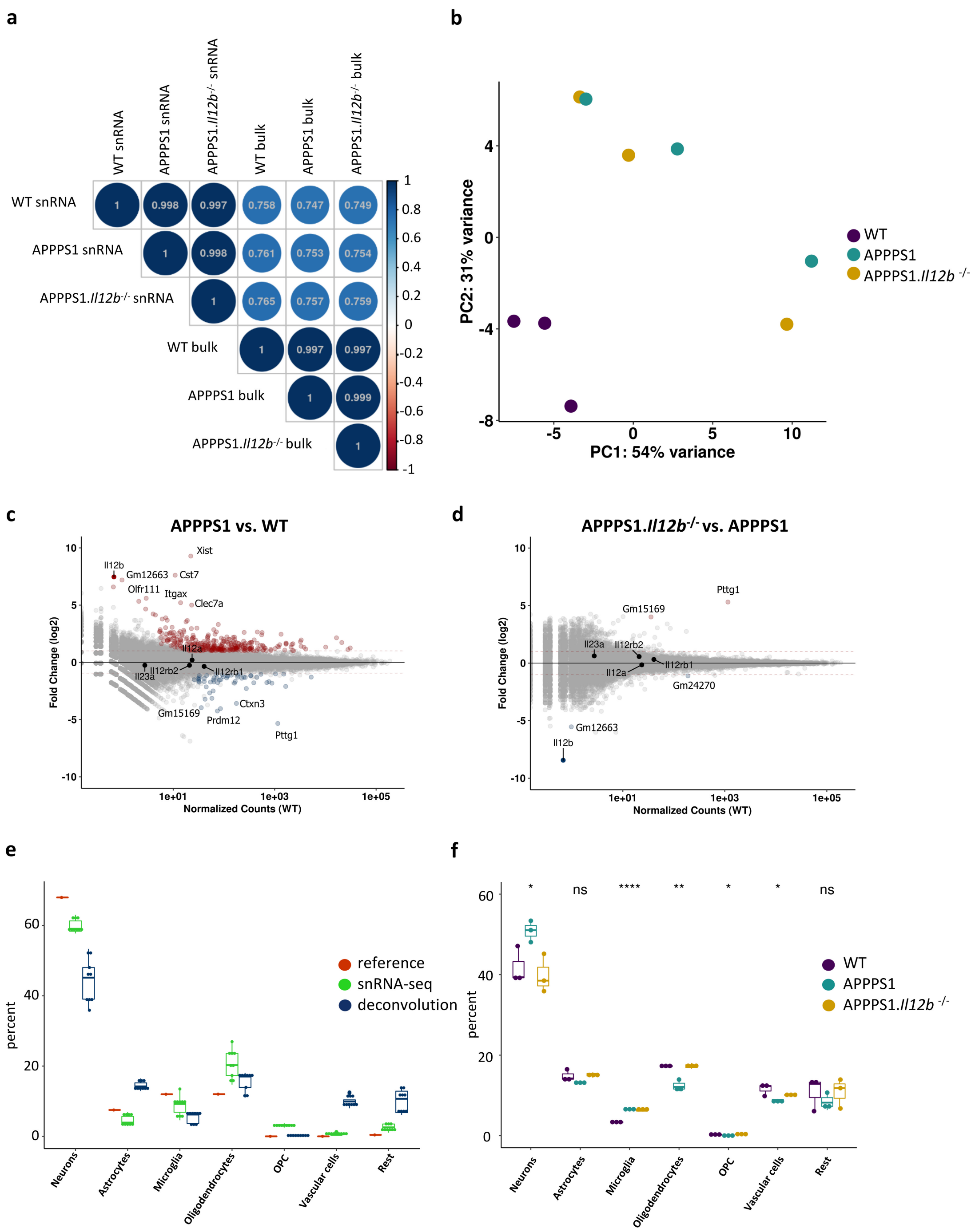


**Extended Data 3: snRNA-seq data from hippocampus correlate well with bulk RNA-seq data and reflects snRNA-seq cellular proportion distribution. a)** snRNA-seq and bulk RNA-seq data showed good correlation with R ≥ 0.75. **b)** Principal component analysis of biological replicates (bulk RNAseq), each replicate represented by one dot. WT datasets cluster together, and as expected APPPS1 and APPPS1.Il12b$^{-/-}$ are more dispersed. **c) – d)** Scatter plots comparing bulk gene expression of **c)** APPPS1 vs. WT and of **d)** APPPS1.Il12b$^{-/-}$ vs. APPPS1. **e)** Comparison of cellular proportions determined by three independent methods, namely the previously published reference data set(red)[20, 21], snRNA-seq (green) and deconvolution of bulk RNA-seq data (dark blue) Important to note is, that the deconvolution matched the other two reference data sets best in OPC, microglia and oligodendrocytes. **f)** Cellular proportions gathered from deconvolved bulk RNA-seq data across all three genotypes showed enhanced number of microglia in APPPS1 and APPPS1.Il12b$^{-/-}$ and a rescue of oligodendrocytes numbers in APPPS1.Il12b$^{-/-}$. * $p \leq 0.05$ (One-way ANOVA, multiple comparison test).

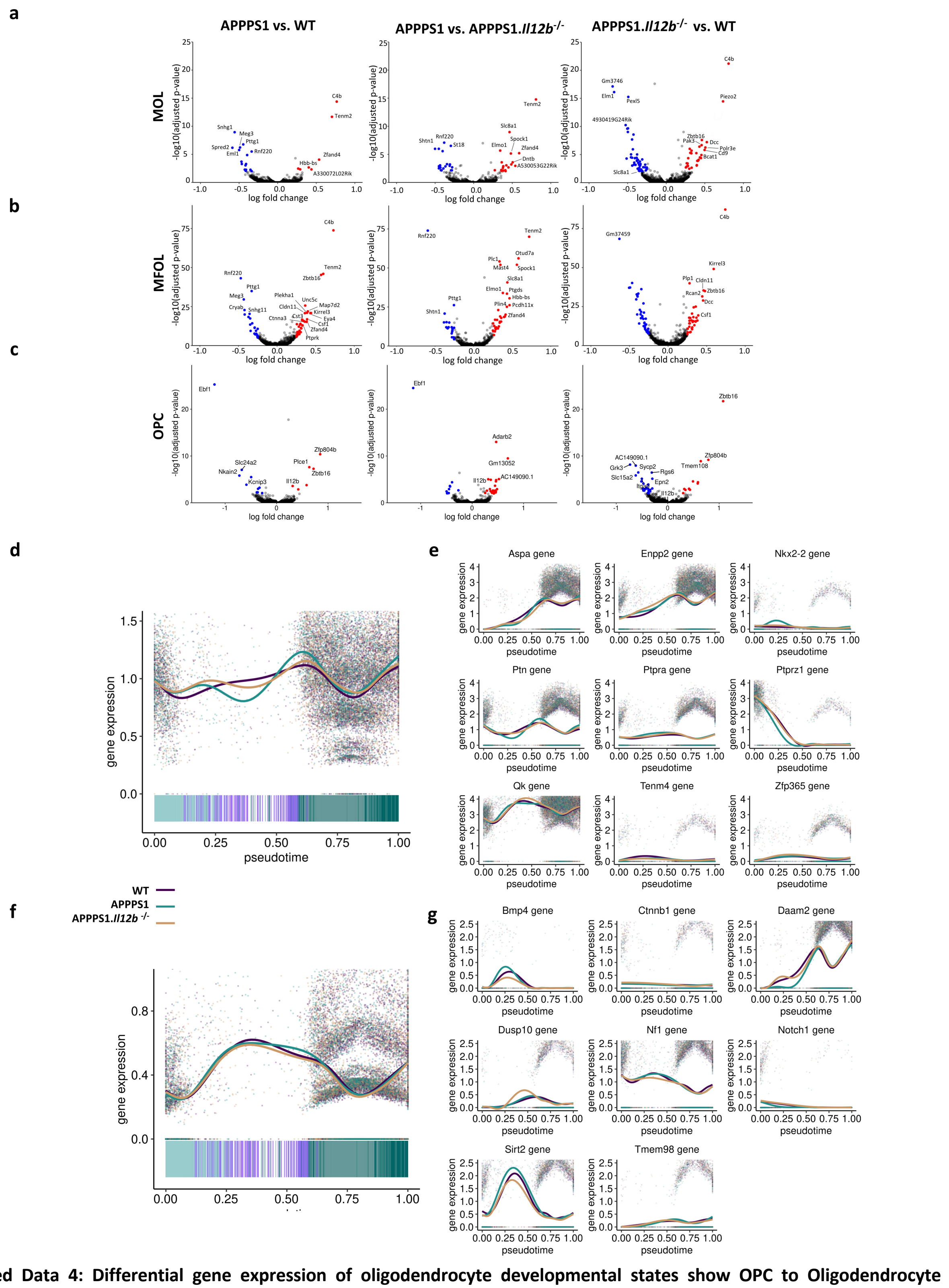


**Extended Data 4: Differential gene expression of oligodendrocyte developmental states show OPC to Oligodendrocyte maturation is unchanged in diseased APPPS1 mice.** Volcanoplots showing differentially regulated genes across all genotypes in **a)** MOL, **b)** MFOL and **c)** OPC. Pseudotime analysis for genes involved in **d) e)** positive regulation (GO:0048714) of oligodendrocyte maturation and **f) g)** negative regulation (GO:0048715) of oligodendrocyte maturation show no strong difference across genotypes.

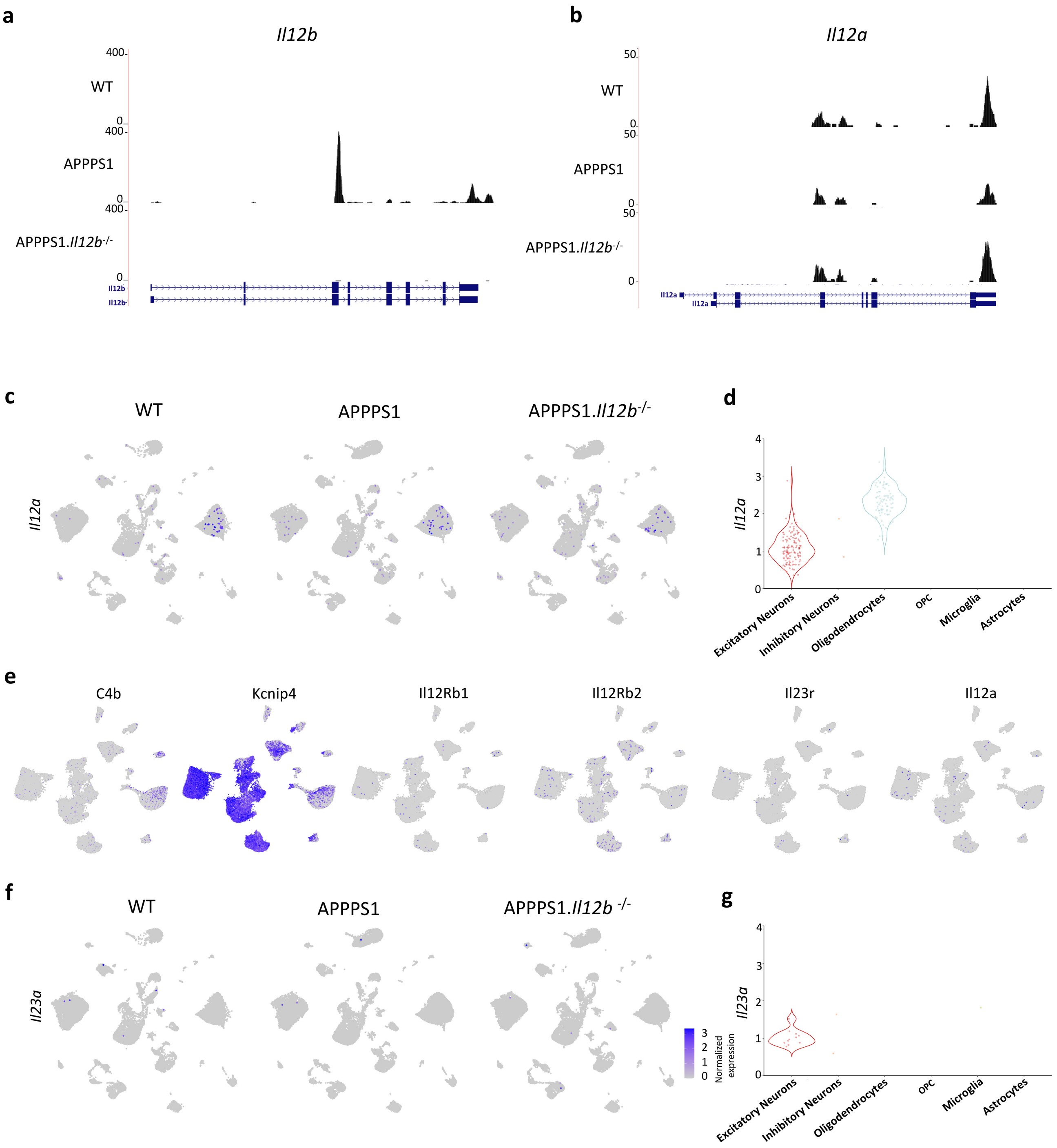


**Extended Data 5: Il12b transcripts are specifically induced in diseased APPPS1 while Il12a is present in all genotypes and Il23a appears almost absent. a)-b)** Genome tracks showing **a)** AD-specificity of Il12b transcripts while **b)** Il12a transcripts are also present in WT and APPPS1.Il12b$^{-/-}$. **c), d)** Il12a is only detectable in excitatory neurons and oligodendrocytes. **e)** snRNA-seq data from Habib and colleagues showing Il12Rb1 expression in Kcnip4 positive cell clusters, marking neurons. Il12Rb2 transcripts are detected in C4b expressing cell cluster, marking oligodendrocytes, as well as Kcnip4 positive cluster. Il23a is almost absent, while Il12a is present in oligodendrocytes and neurons[26]. **f)** Featureplot of own snRNA-seq data as well as **g)** violin-plots show only very few Il23a counts in excitatory neurons.

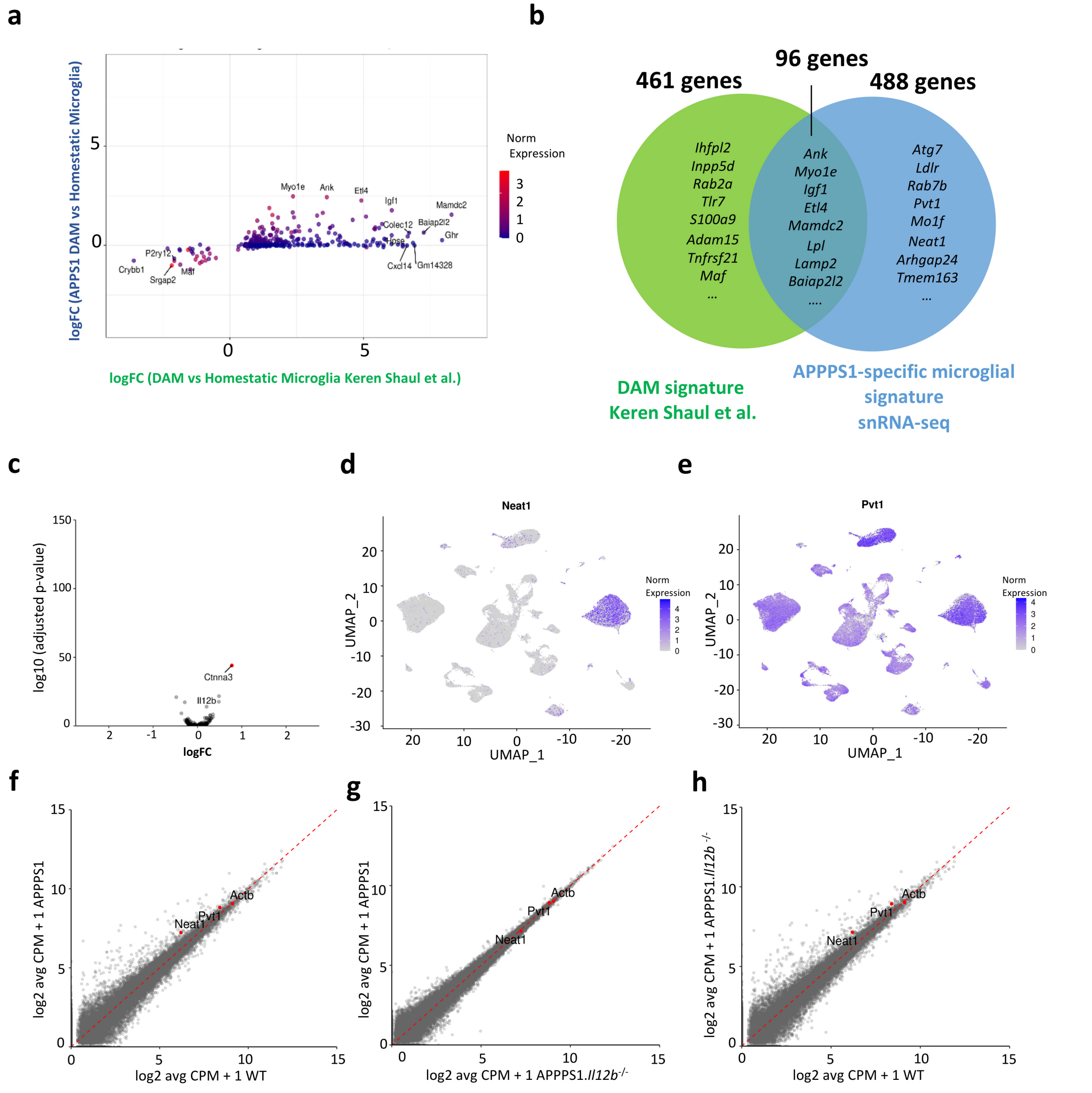


**Extended Data 6: Extended Data 6: Inflammatory protein and non-protein coding gene signature present in both AD-related phenotypes. a)** Gene signature of 5xFAD derived DAM vs. homeostatic microglia27 compared to APPPS1 DAM vs. homeostatic microglia of own snRNA-seq data sets depicted similarities, **b)** Partial overlap of the microglial gene signature of the disease-associated cluster captured in our snRNA-seq data set (blue) with the previously published 5xFAD DAM signature **c)** Volcanoplot comparing microglial gene signature between APPPS1 vs. APPPS1.Il12b-/-. UMAP featureplots showing non-coding transcripts of **d)** Neat1 and **e)** Pvt1. **f)-h)** Volcanoplot comparing **f)** APPPS1 vs. WT **g)** APPPS1 vs. APPPS1.Il12b-/- and **h)** APPPS1.Il12b-/- vs, WT showing that the non coding lncRNA Neat1 and Pvt1 were upregulated in both AD-related phenotype.

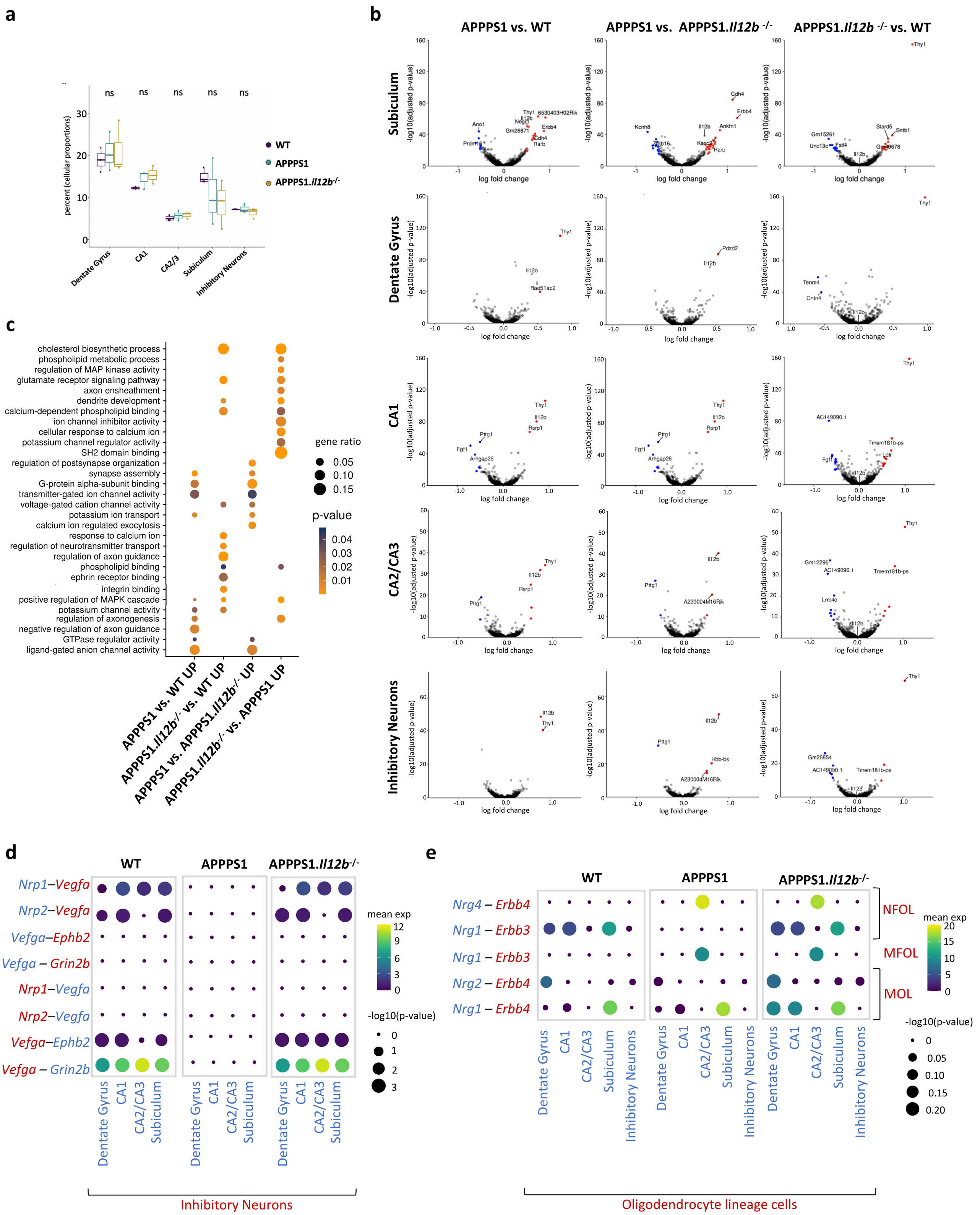


Figure legend on following page

**Extended Data 7: IL-12 signaling leads to transcriptional changes in neurons a)** Subicular neurons showed the strongest decrease of detected nuclei in APPPS1 and APPPS1.Il12b -/- compared to WT. Volcanoplot showing differentially regulated genes across genotypes in **b)** subiculum, dentate gyrus, CA1, CA2/3 and inhibitory neurons. Red and blue dots are determined as statistically significant with p-value ≤ 0.01 and log2 fold-change ± > 0.5 g) GO analysis of genes upregulated in subiculum comparing APPPS1 vs. WT, APPPS1.Il12b-/- vs. WT, APPPS1 vs APPPS1.Il12b-/- and APPPS1.Il12b-/- vs APPPS1. d)-e) Using cellphoneDB35, 36, dot plot showing the predicted receptor ligand interactions with between d) neuronal cell types and e) oligodendrocytes and neurons in WT, APPPS1 and APPPS1.Il12b-/- . p-values are indicated by the circle sizes and means of the average expression level is indicated by the color.
